# Inter-homologue repair in fertilized human eggs?

**DOI:** 10.1101/181255

**Authors:** Dieter Egli, Michael V. Zuccaro, Michal Kosicki, George M. Church, Allan Bradley, Maria Jasin

## Abstract

Many human diseases have an underlying genetic component. The development and application of methods to prevent the inheritance of damaging mutations through the human germline could have significant health benefits, and currently include preimplantation genetic diagnosis and carrier screening. Ma et al. take this a step further by attempting to remove a disease mutation from the human germline through gene editing^1^. They assert the following advances: (i) the correction of a pathogenic gene mutation responsible for hypertrophic cardiomyopathy in human embryos using CRISPR-Cas9 and (ii) the avoidance of mosaicism in edited embryos. In the case of correction, the authors conclude that repair using the homologous chromosome was as or more frequent than mutagenic nonhomologous end-joining (NHEJ). Their conclusion is significant, if validated, because such a “self-repair” mechanism would allow gene correction without the introduction of a repair template. While the authors’ analyses relied on the failure to detect mutant alleles, here we suggest approaches to provide direct evidence for inter-homologue recombination and discuss other events consistent with the data. We also review the biological constraints on inter-homologue recombination in the early embryo.

---

In their first approach, Ma et al. used donor sperm from a patient heterozygous for the *MYBPC3*^***ΔGAGT***^ mutation to fertilize wild-type oocytes, such that half of the embryos started out as wild type at the *MYBPC3* locus and half heterozygous. Fertilized zygotes were injected with Cas9 and an sgRNA directed to create a double-strand break (DSB) in the mutant paternal allele. The authors report that 24% of the embryos at day 3 of development were mosaic, with some cells of the embryo containing the mutant paternal locus, either intact or modified by NHEJ, together with a wild-type locus. Remaining cells of the embryo contained only a detectable wild-type allele. While some zygotes were also co-injected with a wild-type, exogenous, single-stranded oligodeoxynucleotide template (ssODN) with two synonymous mutations, no mutations consistent with ssODN-templated repair were detected. Furthermore, 'wild-type only’ cells were present at a similar frequency both in the presence and absence of the ssODN. The authors infer that these cells arose by homology-directed repair (HDR) of the mutant paternal allele using the wild-type maternal allele as a template, i.e., inter-homologue recombination, leading to gene correction.

In a second approach, earlier, MII-phase oocytes were coinjected with Cas9 complexes and donor sperm. In this case, mosaicism was not detected, except in a single embryo, which contained both 'wild-type only’ cells and ones heterozygous for wild-type and ssODN-templated alleles. Although wild-type embryos were expected at 50% frequency, they appeared to comprise 72% of embryos. The authors argue that the excess (22%) of apparently wild-type embryos arose in the MII-injected oocytes by HDR using the maternal allele to correct the paternal allele, as in the zygote injections, and rarely used the donor template. Thus, a major inference of this article is that a DSB generated by Cas9 in human gametes and zygotes is efficiently repaired by inter-homologue recombination (Fig. 1a). The conclusion that the pathogenic allele can be efficiently corrected without mosaicism has far-reaching implications for the authors’ stated goal of using such methods to address the public health burden of monogenic disease.

**Figure 1.**
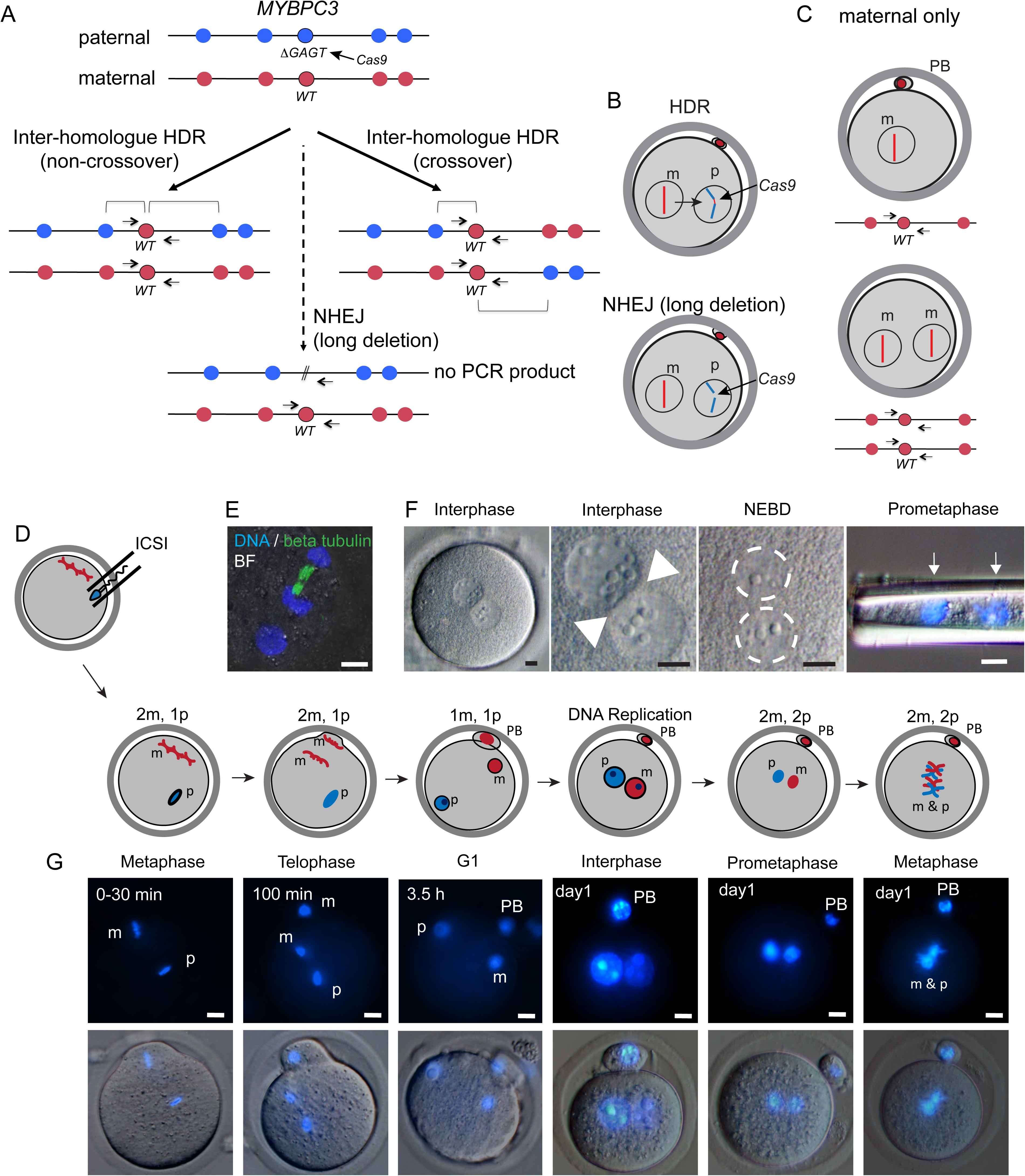
Constraints on gene editing by inter-homologue recombination in the early human embryo. **A.** Possible repair outcomes after a Cas9-induced DSB at the paternal *MYBPC3^ΔGAGT^* locus. Red and blue circles indicate unique maternal and paternal genetic variants. Inter-homologue HDR results in gene conversion of the paternal allele by the wild-type (WT) maternal allele. The repair outcome can be a non-crossover or a crossover. Only one outcome of crossing-over is shown in which the recombined chromosomes are in the same nucleus. The alternative is that the recombined chromosomes segregate to different daughter cells, such that loss of heterozygosity would occur on the chromosome from the point of the HDR event to the telomere in both daughter cells, one with homozygosity for the maternal chromosome and the other for the paternal chromosome. This outcome would be expected in half of the crossing-over events that underwent inter-homologue HDR in G2 phase. NHEJ events are also possible that result in the deletion of a primer-binding site used for genotyping. **B.** Schematic of possible repair outcomes after Cas9 cleavage in the human zygote from panel A. m, maternal chromosome, p, paternal chromosome. **C.** Parthenogenesis after fertilization failure with (top) and without (bottom) second polar body (PB) extrusion. Outcomes of A-C are indistinguishable in genotyping assays using flanking PCR primers alone. **D.** Schematic of intracytoplasmic sperm injection (ICSI) followed by progression through the first cell cycle during day 1 of development. The number of maternal and paternal genomes are indicated at each phase of the cell cycle. **E.** Immunofluorescence of a mouse zygote at telophase of the second maternal meiotic division. Note that only the maternal genomes are attached to microtubules, while the paternal genome begins to form an interphase nuclear membrane to replace the sperm membrane. BF= bright field. **F.** Progression of human zygotes through the first cell cycle from the two-pronuclear stage to prometaphase, when the two genomes can be removed from the egg by a needle. Note the separation of the two genomes (arrows and dashed circles). NEBD, pronuclear envelope breakdown. **G.** Cell cycle progression during day 1 in fertilized mouse zygotes. Of 23 mouse eggs, none showed direct contact between the maternal and paternal genomes. Size bars (independent of color or length in each panel) = 10 μm. Panels one and four in F and panels four to six in G are as published in reference^12^ and in reference^13^.

Given the far-reaching implications, providing direct evidence for correction of the pathogenic allele, rather than the inability to detect the mutant allele, cannot be overemphasized. Another study in mice similarly relied on the absence of a mutant allele to conclude repair by inter-homologue recombination^2^. Considering the data presented in Ma et al., alternatives to recombination between homologues are possible and would seem more likely, as the cell biology of fertilized eggs would appear to preclude the direct interaction between the maternal and paternal genomes required for inter-homologue HDR. Therefore, clear evidence for a novel linkage of maternal and paternal alleles is an imperative for any embryo that would be considered for future implantation.

Novel DNA linkages can be detected directly by sequencing, but the authors do not indicate whether their genome sequencing data was phased to detect the incorporation of the wild-type sequence from one of the maternal homologues at the DSB into the paternal chromosome (Fig. 1A). New parental linkages can also be directly assayed by long-range PCR using allele-specific primers; the only requirement is that SNPs/indels exist to distinguish the maternal and paternal chromosomes in the vicinity of the mutation^3^. Of relevance, this type of analysis can be performed on individual oocytes^4^.

Without direct verification of inter-homologue recombination at the mutant allele, attempts should be made to rule out other types of events. During gene editing, NHEJ is usually considered to lead to small indels at DSB sites. However, with appropriate experimental design, long deletions and other events can be detected in cultured cells and in both mouse and pig zygotes^5-7^. In Ma et al., genotyping involved amplification of a ∼534 bp fragment in which the *MYBPC3*^***Δ**GAGT*^ mutation is ∼200 bp from a primer-binding site. Deletions >200 bp would be sufficient to remove this primer-binding site and lead to amplification only of the maternal allele (Fig. 1A, B), giving the misleading appearance of a corrected paternal allele. To detect longer deletions, a matrix of primer pairs need to be tiled at increasing distances on both sides of the mutation. In a study designed to systematically score these events, Cas9-induced DSBs in mouse embryonic stem cells were found to resolve into large deletions (250-9500 bp) in approximately 20% of edited cells (M.K. and A.B., unpublished results). This approach remains imperfect to detect all events, though, because very large deletions or other events such as translocations prevent amplification and thus escape characterization. Given the ramifications, more studies of this type are required to quantify these events at other loci, particularly in embryos. Because fertilization by mutant sperm in the Ma et al. study can be confidently inferred only for mosaic embryos, this type of analysis is not suitable for embryos derived from MII-phase oocyte injections. Thus, linkage analysis is necessary in these cases.

Are there other possible outcomes that can result in a wild-type genotype in a PCR assay but not involve interhomolog recombination? Zygotes with a single pronucleus are not uncommon after intracytoplasmic sperm injection, occurring in ∼10% of fertilization attempts, and are mostly of parthenogenetic origin, containing only the maternal genome^8^ (Fig. 1C). These zygotes are normally discarded, and Ma et al. show the presence of two pronuclei, although they do not provide information on the number and types of abnormal fertilizations. Furthermore, parthenogenesis can also result in zygotes with two maternal genomes when extrusion of the second polar body fails (Fig. 1C)^9,10^. A paternal contribution was verified by cytogenetic analysis in some of the stem cell lines generated from embryos by Ma et al. (2/6); reporting on the presence of unique paternal polymorphisms in all embryos would address the frequency of parthenogenesis. It also remains possible that a fraction of embryos derived from successful fertilization with mutant sperm are at more risk of paternal chromosome loss due to the occurrence of the Cas9-induced DSB.

Although inter-homologue recombination in fertilized oocytes and zygotes cannot currently be excluded, physical separation of maternal and paternal genomes would be expected to be a substantial impediment. Upon fertilization, distinct maternal and paternal nuclei form (pronuclei), such that the two genomes are separate in a cell that is more than 100 μm in diameter (Fig. 1D-G). This may prevent the incorporation of paternal chromosomes into the oocyte MII spindle (Fig. 1E). During the first interphase, maternal and paternal pronuclei migrate from the site of their formation towards the center of the zygote, but the separation persists throughout interphase (Fig. 1F,G), at which time individual nuclei can be manipulated^11^. In both human and mouse zygotes, maternal and paternal genomes undergo DNA replication in separate nuclei, and enter the first mitosis as separate entities, at which time they can still be manipulated separately (Fig. 1F,G). Merging of maternal and paternal chromosomes does not occur until microtubule action assembles both genomes on a common metaphase plate at the first mitosis ^12,13^. Therefore, direct interactions between maternal and paternal genomes required for inter-homologue repair do not seemingly occur until embryos enter the 2-cell stage when the two genomes are packaged within the same nucleus.

Although the study of DSB repair in human embryos is in its infancy, inter-homologue recombination in mitotic cells appears to be significantly less frequent than inter-sister recombination (or NHEJ), which may be due, at least in part, to the much larger nuclear volume homologues occupy compared to sister chromatids^14^. By contrast, inter-homologue recombination in meiosis, which is essential for the reductional division to form gametes, is efficient, likely due to the large number of DSBs that are programmed to form on each chromosome to promote homologue pairing^15^. It is important to note, however, that meiotic inter-homologue recombination occurs during fetal development in females^16^ and so it is temporally removed from the events described in Ma et al. Whether meiotic recombination factors are still expressed and active in MII-phase oocytes decades later has not been examined as far as we are aware.

In summary, the conclusion of gene correction in human embryos requires further investigation, including direct verification. Efficient inter-homologue recombination in embryos in which the maternal and paternal genomes are undergoing distinct biological programs and in distinct nuclei would be a stunning biological finding. But it would also put cells at risk for unmasking deleterious recessive alleles through loss of heterozygosity (not shown in Fig. 1A). The clinical implications of gene editing in human embryos are substantial. While gene editing could reduce disease-causing alleles, inadvertent changes to the human germline like long deletions and loss of heterozygosity have not been ruled out. Thus, each embryo needs to be carefully evaluated to confirm (or not) gene correction and lack of mosaicism. Despite the inherent limitations imposed on such research, it is essential that conclusions regarding the ability to correct a mutation in human embryos be fully supported. Absent such data, the biomedical community and, critically, patients with disease-causing mutations interested in such research must be made aware that numerous challenges in gene correction remain.

## Author contribution statement

D.E. and M.J. wrote the paper with contributions from M.K., A.B, M.Z. and G.M.C. M.Z. and D.E. performed ICSI and imaging of eggs. M.K. and A.B. provided unpublished information.

## References

1 Ma, H. et al. Correction of a pathogenic gene mutation in human embryos. Nature, doi:10.1038/nature23305 (2017).

2 Wu, Y. et al. Correction of a genetic disease in mouse via use of CRISPR-Cas9. Cell stem cell 13, 659–662, doi:10.1016/j.stem.2013.10.016 (2013).

3 Jeffreys, A. J. & May, C. A. Intense and highly localized gene conversion activity in human meiotic crossover hot spots. Nat Genet 36, 151–156, doi:10.1038/ng1287 (2004).

4 Cole, F. et al. Mouse tetrad analysis provides insights into recombination mechanisms and hotspot evolutionary dynamics. Nat Genet 46, 1072–1080, doi:10.1038/ng.3068 (2014).

5 Shin, H. Y. et al. CRISPR/Cas9 targeting events cause complex deletions and insertions at 17 sites in the mouse genome. Nature communications 8, 15464, doi:10.1038/ncomms15464 (2017).

6 Whitworth, K. M. et al. Use of the CRISPR/Cas9 system to produce genetically engineered pigs from in vitro-derived oocytes and embryos. Biology of reproduction 91, 78, doi:10.1095/biolreprod.114.121723 (2014).

7 Parikh, B. A., Beckman, D. L., Patel, S. J., White, J. M. & Yokoyama, W. M. Detailed phenotypic and molecular analyses of genetically modified mice generated by CRISPR-Cas9-mediated editing. PloS one 10, e0116484, doi:10.1371/journal.pone.0116484 (2015).

8 Sultan, K. M., Munne, S., Palermo, G. D., Alikani, M. & Cohen, J. Chromosomal status of uni-pronuclear human zygotes following in-vitro fertilization and intracytoplasmic sperm injection. Human reproduction (Oxford, England) 10, 132–136 (1995).

9 Kim, K. et al. Recombination signatures distinguish embryonic stem cells derived by parthenogenesis and somatic cell nuclear transfer. Cell stem cell 1, 346–352 (2007).

10 Paull, D. et al. Nuclear genome transfer in human oocytes eliminates mitochondrial DNA variants. Nature 493, 632–637, doi:nature11800 [pii] 10.1038/nature11800 [doi] (2013).

11 Kattera, S. & Chen, C. Normal birth after microsurgical enucleation of tripronuclear human zygotes: case report. Human reproduction (Oxford, England) 18, 1319–1322 (2003).

12 Egli, D. et al. Reprogramming within hours following nuclear transfer into mouse but not human zygotes. Nature communications 2, 488, doi:ncomms1503 [pii] 10.1038/ncomms1503 [doi] (2011).

13 Egli, D., Rosains, J., Birkhoff, G. & Eggan, K. Developmental reprogramming after chromosome transfer into mitotic mouse zygotes. Nature 447, 679–685 (2007).

14 Stark, J. M. & Jasin, M. Extensive loss of heterozygosity is suppressed during homologous repair of chromosomal breaks. Mol Cell Biol 23, 733–743 (2003).

15 Kauppi, L. et al. Numerical constraints and feedback control of double-strand breaks in mouse meiosis. Genes Dev 27, 873–886, doi:10.1101/gad.213652.113 (2013).

16 Baker, T. G. A QUANTITATIVE AND CYTOLOGICAL STUDY OF GERM CELLS IN HUMAN OVARIES. Proceedings of the Royal Society of London. Series B, Biological sciences 158, 417–433 (1963).

